# Behavioral and Hematological Response of *Clarias gariepinus* juvenile exposed to Acute concentrations of Aqueous Extract of Siam Weed (*Chloromolaena odorata*) leaf as Anesthetic agent

**DOI:** 10.1101/181164

**Authors:** Iheanacho Stanley Chidi, Nworu Shedrack

**Affiliations:** Department of Fisheries and Aquaculture, Federal University Ndufu Alike Ikwo, Ebonyi State, Nigeria; Department of Fisheries and Aquaculture, Ebonyi State University Abakaliki, Ebonyi State, Nigeria

**Keywords:** Effects, *C. odorata*, Behavior, Hematology, *C. gariepinus*

## Abstract

This experiment was carried out to investigate the effect of Siam Weed (*chloromelena odorata*) on the heamatology of *Clarias gariepinus* juvenile. A total of one hundred and fifty (150) juvenile of *Clarias gariepinus* were randomly assigned to different concentrations of *C. odorata* leave aqueous extract in a completely randomize design (CRD). The concentrations were 50mg/l, 100mg/l, 150mg/l, 200mg/l. Distilled water (0.00 mg/l) was used as the control. The fish exhibited stressful behavior which was higher as the concentration of *Chromolaena odorata* leave extract increased. There was a gradual decrease with time until a state of calmness, which was subsequently followed by death. The effect on 96hr exposed period was recorded and blood samples collected at 24hr and 96hr interval. Result on hematological parameters revealed significant difference (P<0.05) among treatments with increase in exposure time for all the blood parameters. *C. odorata* at increased concentrations affected the behavior and hematology of *C. gariepinus.*

## Introduction

Aquaculture is the fastest growing food producing sector in the world (Iheanacho et al. 2017) and has grown exponentially over the years especially in developing countries (James et al. 2001). Over the years, Aquaculture sector has encountered serious setbacks such as mortalities emanating from disease infections, mechanical injuries during transportation stress, sampling and sorting of fish (Romero et al. 2012). Anaesthetics are used with increasing frequency in aquaculture, mainly to reduce the stress and to prevent mechanical damage to fish during handling. Their use is particularly common in stripping, marking, biometry, tagging, artificial reproduction procedures and surgery, thus reducing stress-induced problems such as reduction in feeding and immune function (Ross and Ross 1999; Kolarova et al. 2007; Altun et al. 2009; Yildiz et al. 2013, Opiyo et al. 2013). The need to handle fish without impairing their health or commercial value has led to the development of many techniques to anaesthetize fish (Yildiz et al. 2013). Different anaesthetics act with various intensity driving fish into general anaesthesia, resulting in loss of consciousness, inhibition of reflex activity, and reduction in skeletal muscle tone (Hajek et al. 2006). Quick induction and recovery from anaesthesia is desirable in most cases (Marking and Meyer 1985; Stoskopf 1993).

However, long recovery time is desirable while collecting fish from the wild or where fish must be handled for a longer time in the laboratory. An ideal anaesthetic should possess several attributes such as; non-toxic, inexpensive, simple to administer and result in rapid induction and calm recovery (Pawar et al. 2011). It is of paramount importance to identify the effective dose of anaesthetic for specific fish species since response to the same anaesthetic vary depending on the concentration used and the species of fish (Tyler and Hawkins 1981; Pawar et al. 2011). Common fish anaesthetics include clove oil, sodium bicarbonate, carbon dioxide gas, metomidate, benzocaine, tricaine, methanesulphonate (MS-222), 2-phenoxylethanol and quinaldine (Massee et al. 1995; Palić et al. 2006). Regardless of the agent, the process of anaesthesia in fish develops in a similar way and runs in a progressive pattern.

Fish anaesthetics have positive effects on fish during transportation (Berka 1986), and handling or during surgical procedures (Marking and Meyer 1985). Selecting a suitable anesthetic depends mainly on its effectiveness in immobilizing fish with good recovery rates (Berka et al. 1986). The efficacy of aqueous extracts of some plants such as avocado pear as anaesthetic was been reported by Adebayo et al. (2010). Clove oil was used as anaesthetic on different species of fish (Taylor and Roberts 1999). Similarly, the effectiveness of mistletoe extract as anaesthetic was reported by Popoola et al. (2008).

In recent years there is preference for safe and environmentally friendly piscicides of plant origin than synthetic piscicides for catching fish and clearing pond. This is because ichthvotoxins are less expensive, biodegradable, readily available, easy to handle and save for mankind and the environment (Singh et al. 2010). The deliberate introduction of these plants extract in the aquatic ecosystems could eventually lead to physiological stress in aquatic organisms and ultimately reduction in aquatic productivity (Olufayo 2009). Plants parts have been shown to cause death of fish and changes in biochemical response of *Channa punctatus* (Tiwari and Singh 2004), haematological and histopathological effects on *Clarias gariepinus* (Omoniyi et al. 2002). Ubaha et al. (2012) reported decreased haemoglobin, haematocrit and erythrocytes when they studied the effect of *Hypoestes forskalei* leaf extract on the behavior of *C. gariepinus*. Ojutiku et al. (2013) studied the effect of acute concentration of cypermethrin on juveniles of *C. gariepinus* and reported that white blood cells (WBC), MCV, MCH, PCV and neutrophil levels increased, while RBC and lymphocytes reduced. Although anaesthetia of fish have positive effects on the fish during transportation and handling, some anaesthesias can pose dangerous problems to the fish organs and the blood parameters (Nicula et al. 2010). Knowledge of the haematological characteristics is an important tool that can be used as an effective and sensitive index to monitor physiological and pathological changes in fishes (Iheanacho et al. 2017).

*Chromolaena odorata* is a species of flowering shrub in the sunflower family, Asteraceae. It is native to North America, from Florida and Texasto Mexico and the Caribbean, and has been introduced to tropical Asia, west Africa, and parts of Australia. Common names include Siam Weed, Christmas Bush, and Common Floss Flower. It is sometimes grown as a medicinal and ornamental plant. It is used as a traditional medicine in Indonesia. The weed is used as herbal medicine (Ling et al. 2007) and produces characteristic smell when crushed. *C. odorata* have a complex mixture of flavonoid compounds including aurone, flavones and flavonol (Vital 2008). The flavonoid compounds possess antibacterial, antifungal and antiprotozoal properties (Vital and Rivera 2009).

Siamweed extract (SWE) accelerates hemostasis and wound healing (Phan et al. 2001). The presence of flavonoid an active chemical in *C. odorata* has been confirmed (Vital 2008).This compound is a potent antioxidant (Taleb-contini 2006).

This present study would form a baseline data for assessment of health status as per hematological indices of *C. gariepinus* exposed to acute concentration of aqueous extract of Siam weed (*C. odorata*) leaves under laboratory conditions.

## Materials and Methods

### Area of the Study

The study site was Federal University Ndufu Alike Ikwo, Ebonyi State. Ebonyi State is located approximately within latitude 6°20’N and longitude 8°06’E in the derived savannah of south-Eastern part of Nigeria at an elevation of 117m. The rainfall pattern is bimodal (April-July and September-November) with a short spell in August referred to as August break and annual rainfall of about 1,800-2,000mm. The average temperature is between 25°C in January, 34°C in June and 30°C in November.

### Procurement of the fish

*C. gariepinus* juveniles weighing 50.02 g and total body length of 12.5 cm were procured from a reputable commercial fish Farm in Abakaliki, Ebonyi State. Fish were transported (15-25minutes) with 50 litres gallon filled with fresh water from the farm, to wet laboratory and stocked in 200 litres capacity plastic tank. The fish were acclimatized to laboratory conditions (25°C) for 10 days prior to the experiment. During the acclimation period the fish were fed twice daily commercial fish feed (Coppens, 4mm). Feeding of fish stopped two days prior to the start of the experiment.

### Collection, Indentification and Preparation of the plant

*Chromolaena odorata* leaves sample was collected from the wild at Abakaliki and was identified by a botanist from the Department of Agriculture, Federal University Ndufu Alike Ikwo, Ebonyi State, Nigeria (Plate 1). Samples of *Chromolaena odorata* leaves were washed and shade-dried. It was then pulverised using mechanical blender. Precisely 100 grams of the fine powdered *C. odorata* was weighed using a weighing balance. The weighed sample was soaked in 200mls of distilled water contained in a conical flask and swirled. After 48hours, with interval stirring, the mixture was filtered using Whatman No.1 filter paper into a clean beaker. Extracts obtained was filtered with a membrane filter of pore size 0.45ul to obtain a sterile extract, the filtered extract was centrifuged using centrifugal machine and stored in an air-tight bottle at 4°C.

### Range-Finding

A preliminary (range-finding) test as describe by Rahman et al. (2005) was conducted to determine the main experimental concentrations for the *C. odorata* leaves extract. The main experimental concentrations for the extract were determined based on 0-100% of *C. gariepinus* in 24hours.

### Experimental design

Total of 150 *Clarias gariepinus* juvenile were randomly assigned to five treatments with each treatment containing ten fish. Each treatment was replicated three times in a completely randomized design (CRD). A total of 15 plastic aquaria tanks (2m x 1.5m x 1m) were used for the experiment. Concentrations of 0.00, 50, 100, 150 and 200ml/l of aqueous extracts of *C. odorata* known as treatments were prepared by weighing the aqueous extract and mixing it with cold water. Temperature and pH were determined at the start of the experiment and maintained optimal levels. The fish were exposed to 0.4 g (96 hr LC50) for 4 days (Okorie et al.1992) of the aqueous extract of *Chromolaena odorata*. A set of 10 fish were also simultaneously maintained in distilled water (0.00 mg/l) as the control each time the test was repeated.

### Collection of blood for Analysis

At the end of day 24 and 96hr, two fish from each treatment were randomly taken for collection of blood. Holding the fish firmly, the operculum was lifted and blood collected by puncturing the cardiac into EDTA containers for determining haematological parameters.

### Haematological Analysis

Blood parameters such red blood count (RBC), White blood cell (WBC), Hemoglobin content (Hb) and Pack cell volume (PCV) were determined following the procedure of Dacie and Lewis (2011).

### Water quality parameters

Water quality parameters such as temperature, dissolved oxygen and pH of the experimental tank water were determined using water quality kit (Pro. Kit, Flourida).

### Statistical Analysis

The probit method of Finney (1971) was applied to estimate the 96 hour LC50. Results were reported as mean ± standard error (SE) where appropriate. Mean values were compared with one-way analysis of variance (ANOVA) and considerable variations amongst sets were determined by Ducan multiple range test using SPSS for windows version 20. The degree of significant was set at P<0.05.

## Result

### Water parameter

Mean values of water quality parameters for the different concentrations of *C. odorata* leaves extract and control media to which the test fish *C. gariepinus* were exposed over the 96hour exposure period are presented in Table 1. Mean values of the water temperature were not significantly (*P□*0.05) affected by the concentrations of *C. odorata* leaves extract. On the other hand, pH and dissolved oxygen significantly (*P*<0.05) decreased as the concentrations of *C. odorata* leaves extract increased.

**Table 1:** Mean values of water Quality parameters recorded.

| Parameter | T0 (Control) | T1 (50ml/l) | T2 (100ml/l) | T3 (150ml/l) | T4 (200ml/l) |
| --- | --- | --- | --- | --- | --- |
| Temperature | 28.00 | 28.00 | 28.00 | 28.00 | 28.00 |
| (°C) |  |  |  |  |  |
| DO (mg/l) | 8.70 | 8.10 | 6.90 | 5.50 | 5.10 |
| pH | 6.80 | 6.50 | 6.20 | 6.00 | 5.80 |

### Toxicity Bioassay (Mortality Response)

Mortality in the three replicate of *C. odorata* leave extract concentrations at 96hr period varied significantly (P<0.05) in all the treatments and increased with increase in concentration (Table 2). Mucos was copiously observed on the gills of the dead fish in all the treatments except the control which recorded no mortality. The Probit mortality data (Table 3) shows the mortality and time for 50% mortality (LC50).

**Table 2.**
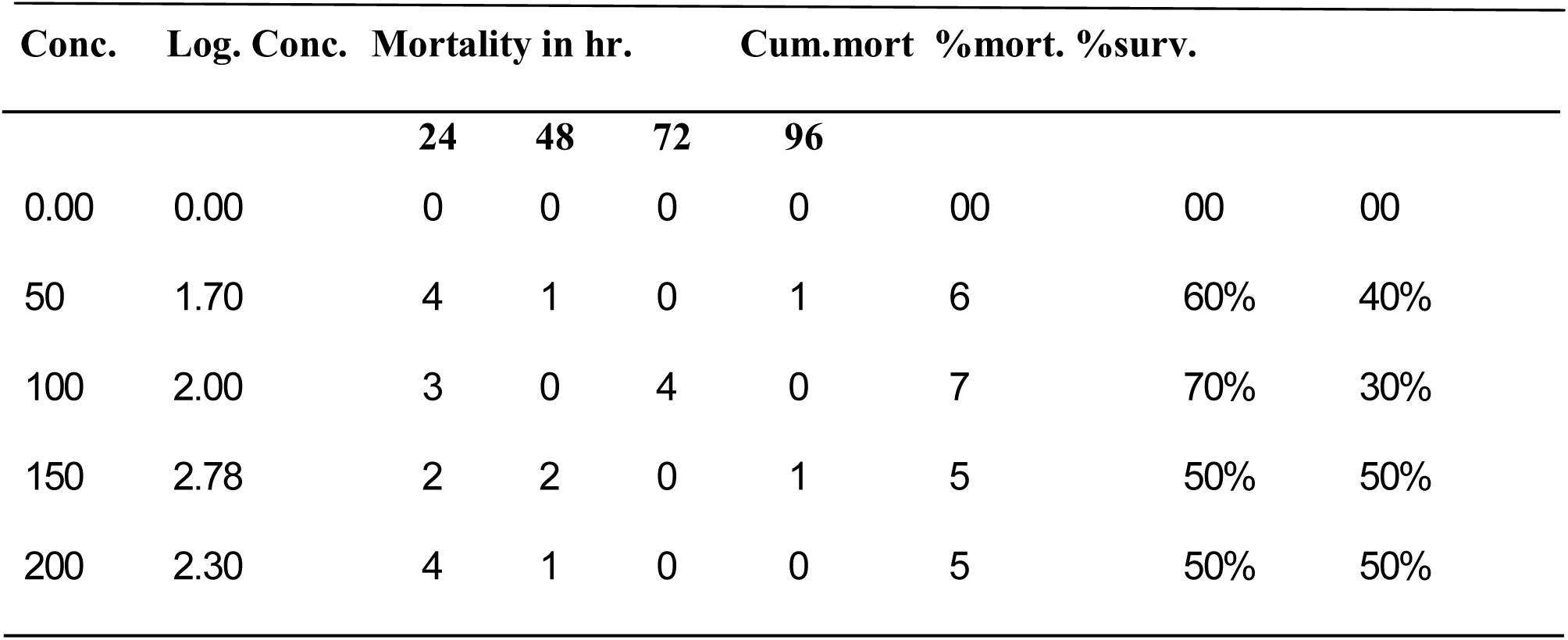
Mortality record for *C. gariepinus* juvenile Exposed to Different Concentration of *C. odorata* leave Extract.

| Conc. | Log. Conc. | Mortality in hr. |  |  | Cum.mort | %mort. | %surv. |
| --- | --- | --- | --- | --- | --- | --- | --- |
|  |  | 24 | 48 | 72 | 96 |  |  |

|  |  |  |  |  |  |  |  |  |
| --- | --- | --- | --- | --- | --- | --- | --- | --- |
| 0.00 | 0.00 | 0 | 0 | 0 | 0 | 00 | 00 | 00 |
| 50 | 1.70 | 4 | 1 | 0 | 1 | 6 | 60% | 40% |
| 100 | 2.00 | 3 | 0 | 4 | 0 | 7 | 70% | 30% |
| 150 | 2.78 | 2 | 2 | 0 | 1 | 5 | 50% | 50% |
| 200 | 2.30 | 4 | 1 | 0 | 0 | 5 | 50% | 50% |

**Table 3.**
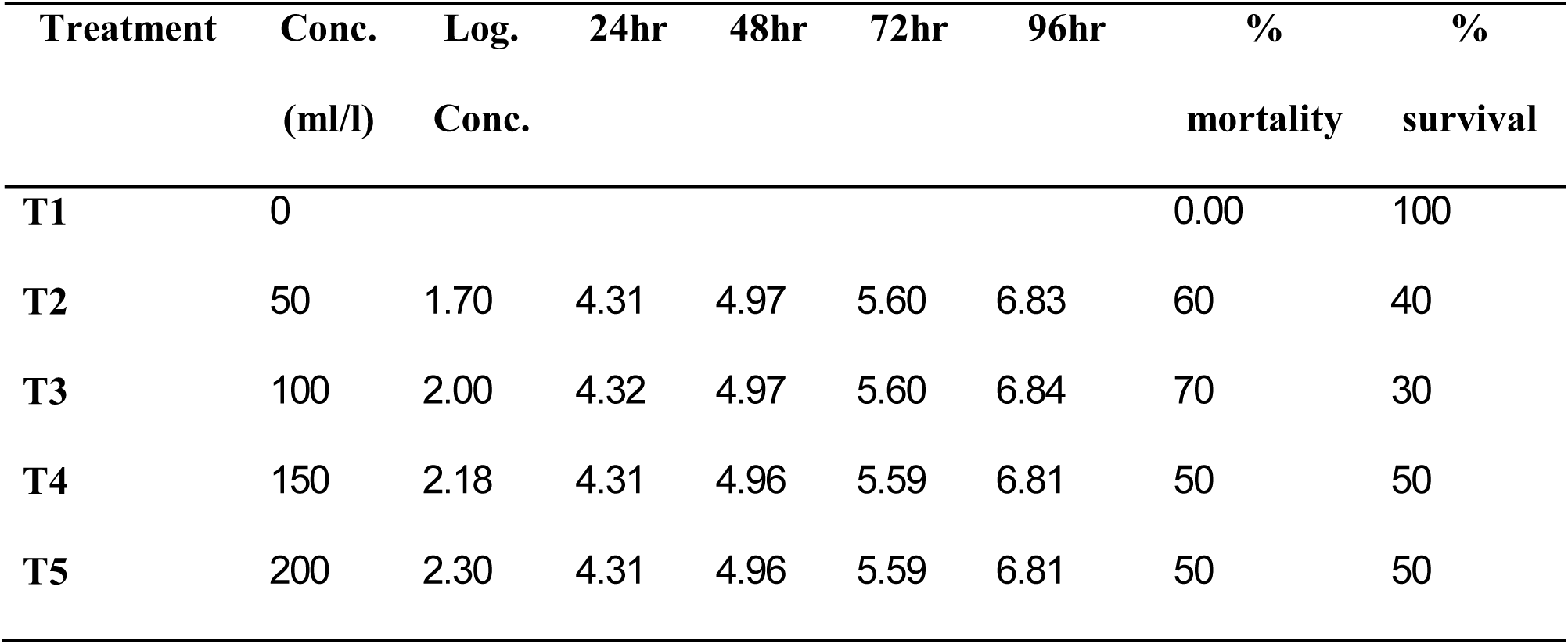
Probit kill for *Clarias gariepinus* juvenile exposed to aqueous extract of *C. odorata.*

### Haematological analysis

Mean values for haematological examination of *C. gariepinus* exposed to acute concentration of aqueous extract of *C. odorata* leaf are presented in the respective figures (Fig.1,2,3 and 4). Result on hematological examination showed significant difference ((*P*<0.05) for all the parameters measured, when compared to the control.

**Fig. 1:**
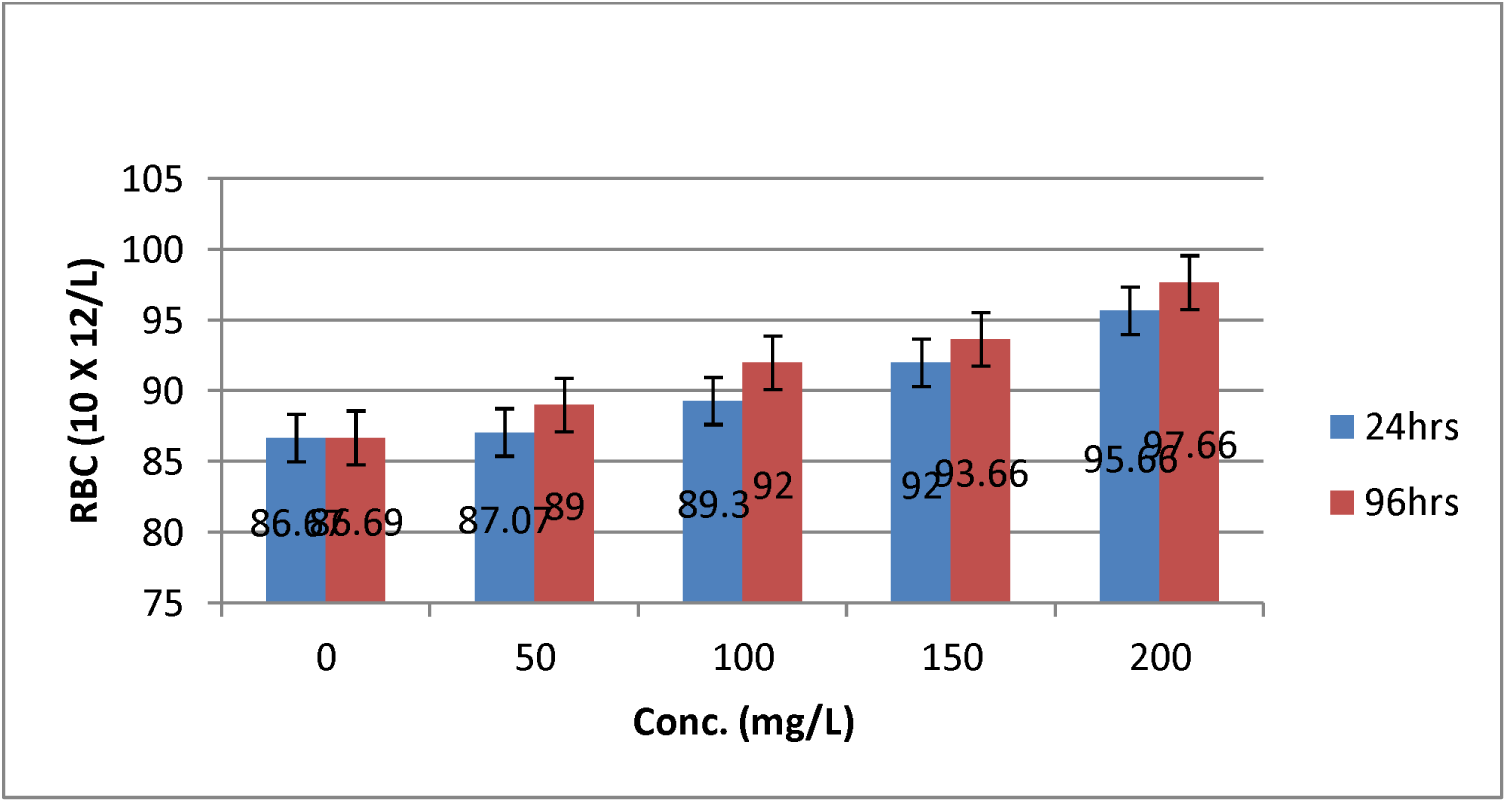
Mean values of Red blood count (RBC) of *C. gariepnus* exposed to acute concentrations of *C. odornata* between 24 and 96hrs.

**Fig. 2:**
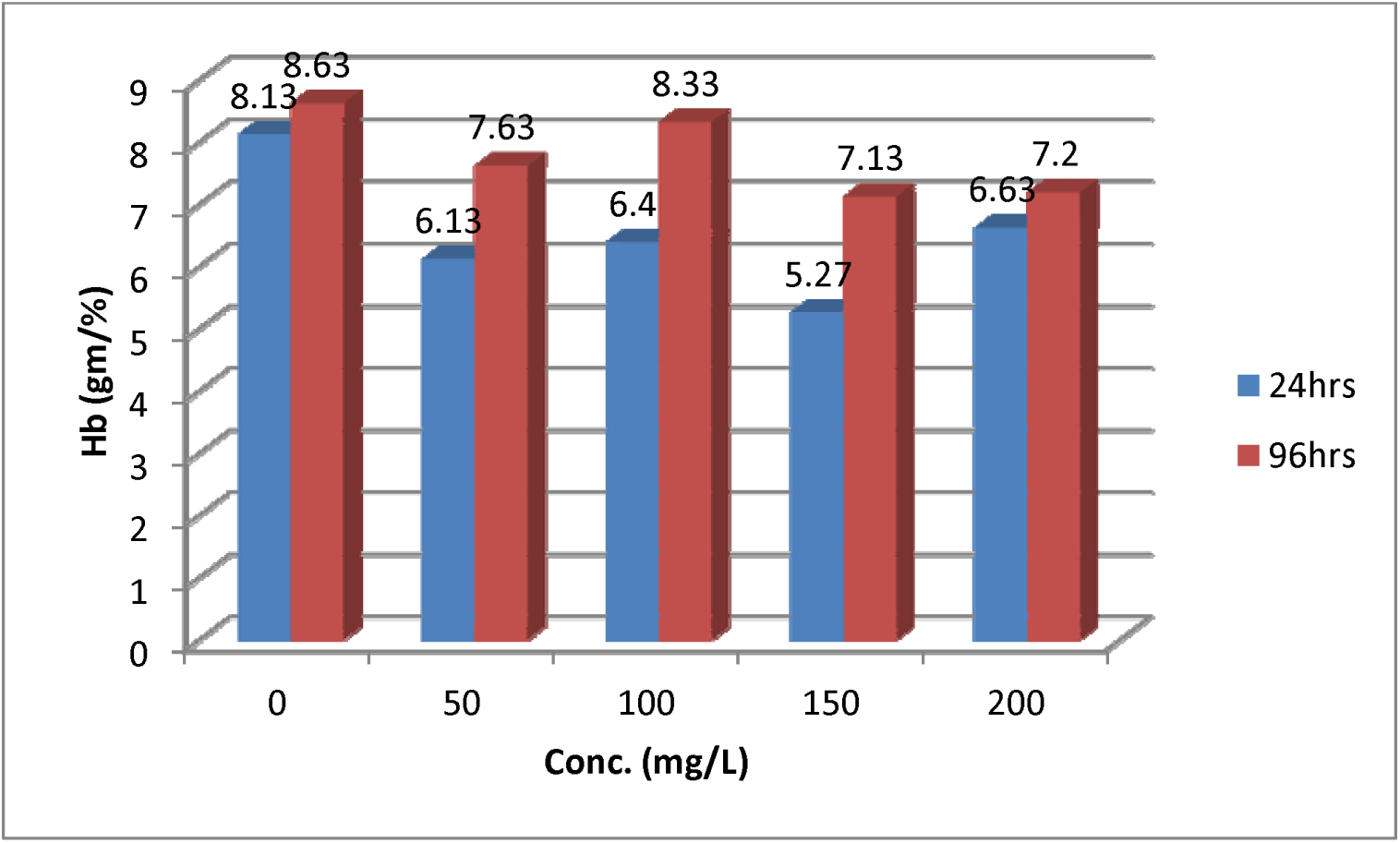
Mean values of Hemoglobin content (Hb) of *C. gariepnus* exposed to acute concentrations of *C. odornata* between 24 and 96hrs

**Fig. 3:**
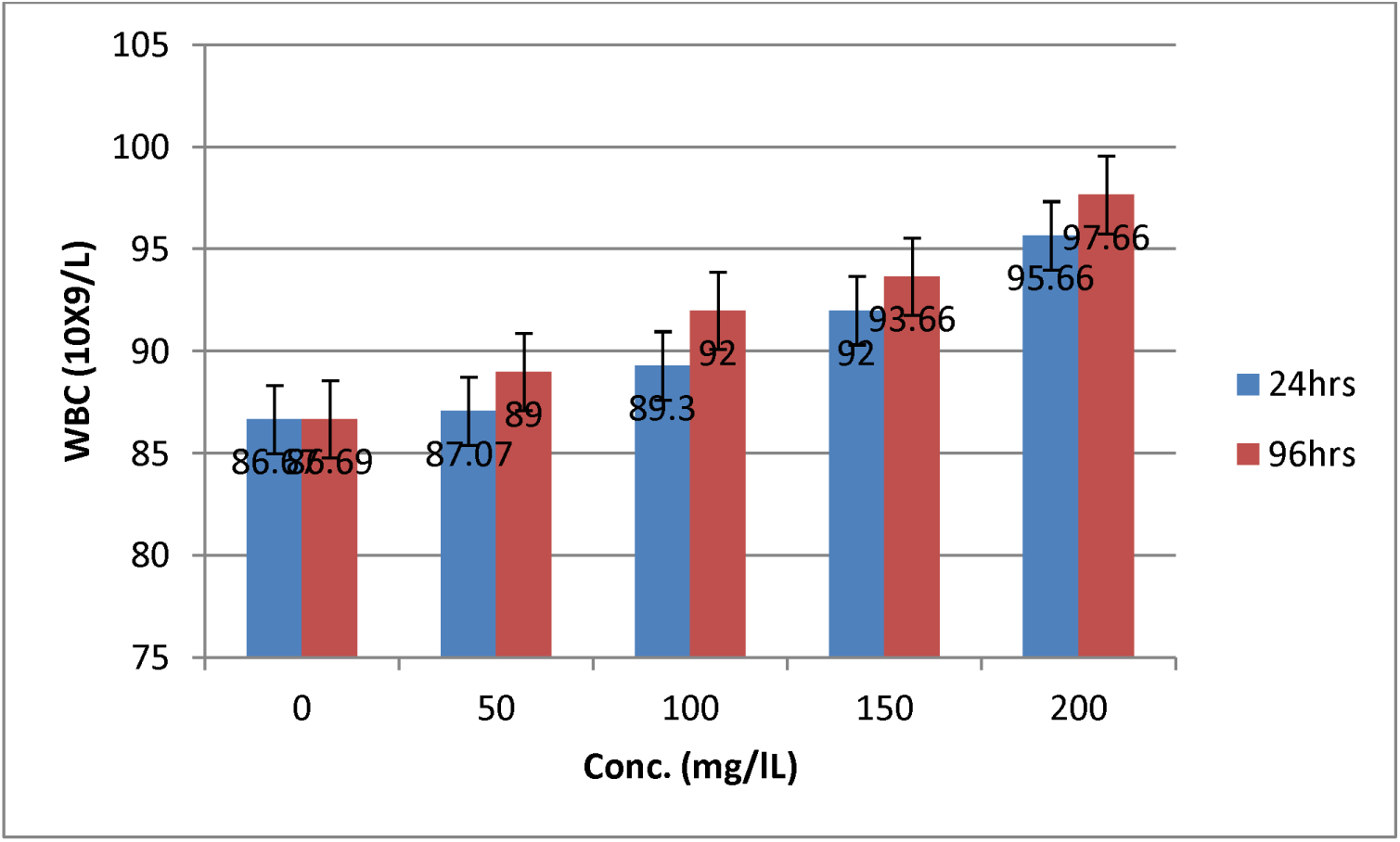
Mean values of White blood cell (WBC) of *C. gariepnus* exposed to acute concentrations of *C. odornata* between 24 and 96hrs

**Fig. 4:**
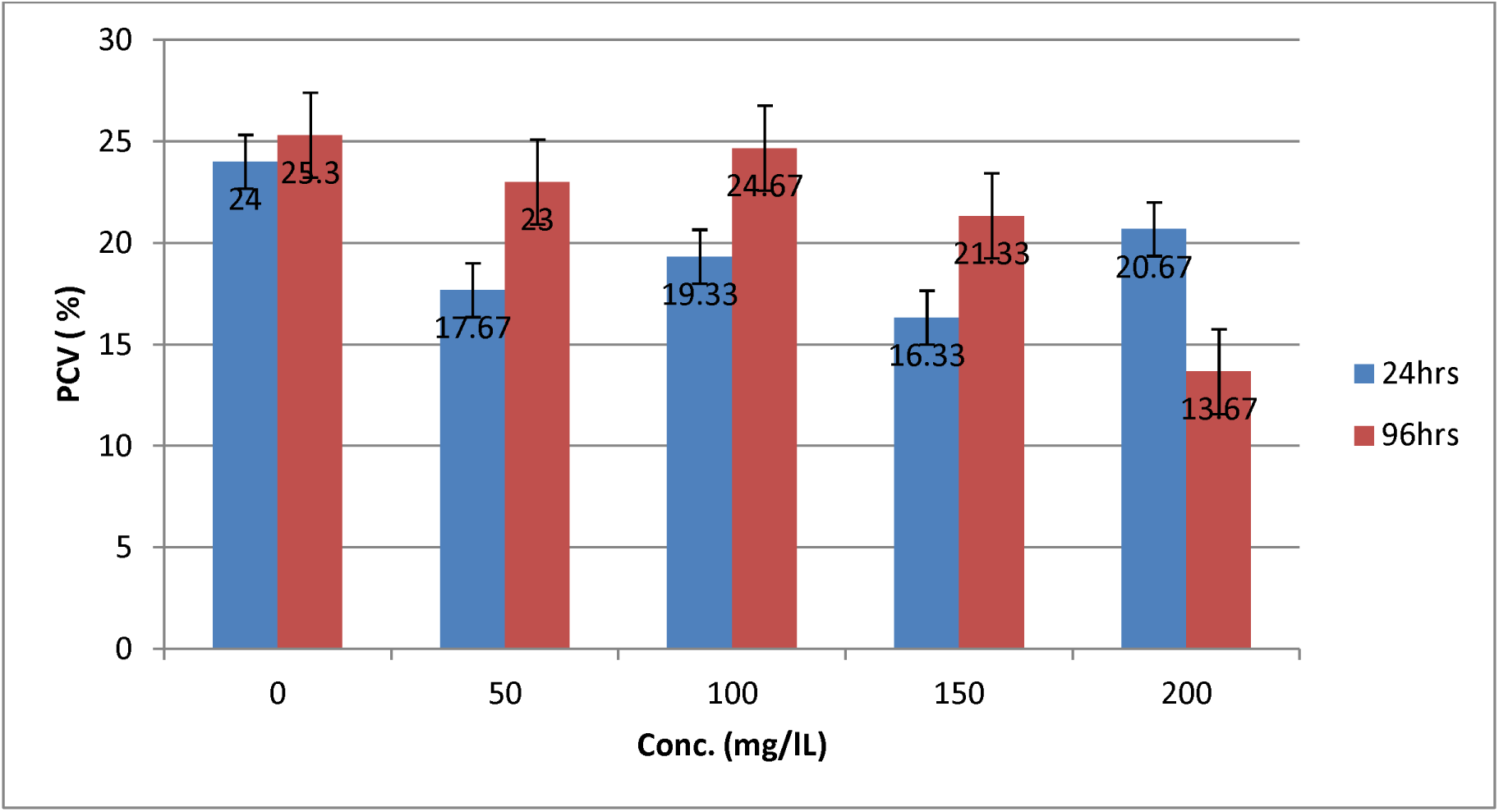
Mean values Pack cell volume (PCV) of *C. gariepnus* exposed to acute concentrations of *C. odornata* between 24 and 96hrs Plate 1: The leaves of Siam weed *Chloromolaena odorata*

## Discussions

### Water Quality parameters

Based on the present findings, changes in the water quality parameters showed that the *C. odorata* concentrations significantly affected the water quality especially dissolve oxygen and pH. Omoniyi et al. (2002), reported a decreased in water quality parameter values when *Clarias gariepinus* was exposed to sub lethal concentrations of tubbaco leaf extract. The variation in the reported result of monitored parameters could be associated with the exposure period and the level of *C. odorata* extract concentrations. The decline in pH with time could be due to the production of acidic metabolites (Haylor 1991). The acidic condition of the experimental tank water may have resulted to a decrease in the level of dissolved oxygen.

### Behavioural Characteristics

The fish exhibited stressful behavior which was higher as the concentration of the solution increased. There was a gradual decrease with time until a state of calmness, which was subsequently followed by death. A similar pattern was reported by Fafioye et al. (2001) on *Clarias gariepinus,* and Rahman et al. (2002) for *Clarias punctatus*. The observed restlessness and uncoordinated swimming in bioassay media might be due to the effect chemical components *C. odorata* leave extract. Akinbulumo (2005) reported that the fish showed a toxic reaction to *Derris elliptica* root powder by surfacing jaws and becoming stupefied.

### Heamatological Parameter

Haematological indices are essential health indicators that reveal the health status of fish before and after an experiment (Iheanacho et al. 2017). The results indicated that PCV values reduced significantly (p□0.05) with increased in concentration, although a mixed trend was observed for 24hrs hematological examination while a corresponding decrease in PCV values was observed for 96hrs compared to the control (Fig.4). Haemoglobin values for treated fish reduced significantly (p□0.05) for both 24hrs and 96hrs with respect to the control (Fig. 2). It was also observed that leucocytes counts increased significantly (P<0.05) (Fig. 3) while erythrocytes counts reduced significantly (*P* < 0.05) with increasing concentrations of Siam weed treated fish groups compared to the control for both periods (24hrs and 96hrs) (Fig. 1). Plants parts (Leaves root and bark) have been shown to cause death of fish and changes in biochemical response of *Channa punctatus* (Tiwari and Singh 2004), haematological and histopathological effects on *Clarias gariepinus* (Omoniyi *et al.,* 2002). The effect of sublethal concentration of formothion has been reported by Singh and Srivastava (1999), with a significant decrease in haematocrit and haemoglobin concentration in *Heteropneustes fossil.* Ubaha *et al.* (2012) where gradual decrease in haemoglobin, pack cell volume and red blood cell count were observed when they studied the effect of *Hypoestes forskalei* leaf extract on the behavior of *C. gariepinus.* The significant increase in the values of white blood cell as the concentration of *C. odorata* leave extract increased could be attributed to the increase in leucocytes synthesis as a defense mechanism against the destruction of erythrocytes. Lymphocytes are the most numerous cells comprising the leucocytes, which function in the production of antibodies and chemical substances serving as a defense against infection (Iheanacho et al. 2017). Ojutiku et al. (2013) studied the effect of acute concentration of cypermethrin on juveniles of *C. gariepinus* and reported that white blood cells (WBC), MCV, MCH, PCV and neutrophil levels increased, while RBC and lymphocytes reduced.

### Conclusion

Although the efficacy of plant materials in aquaculture have been effectively demonstrated, there is need to investigate their effects on fish health. Following the findings of the current study, we conclude that Siam weed (*C. odorata*) altered hematological parameters and affected negatively, the behavior of *Clarias gariepinus.*

## Acknowledgement

Authors are grateful to family members

---

**Plate 1:**
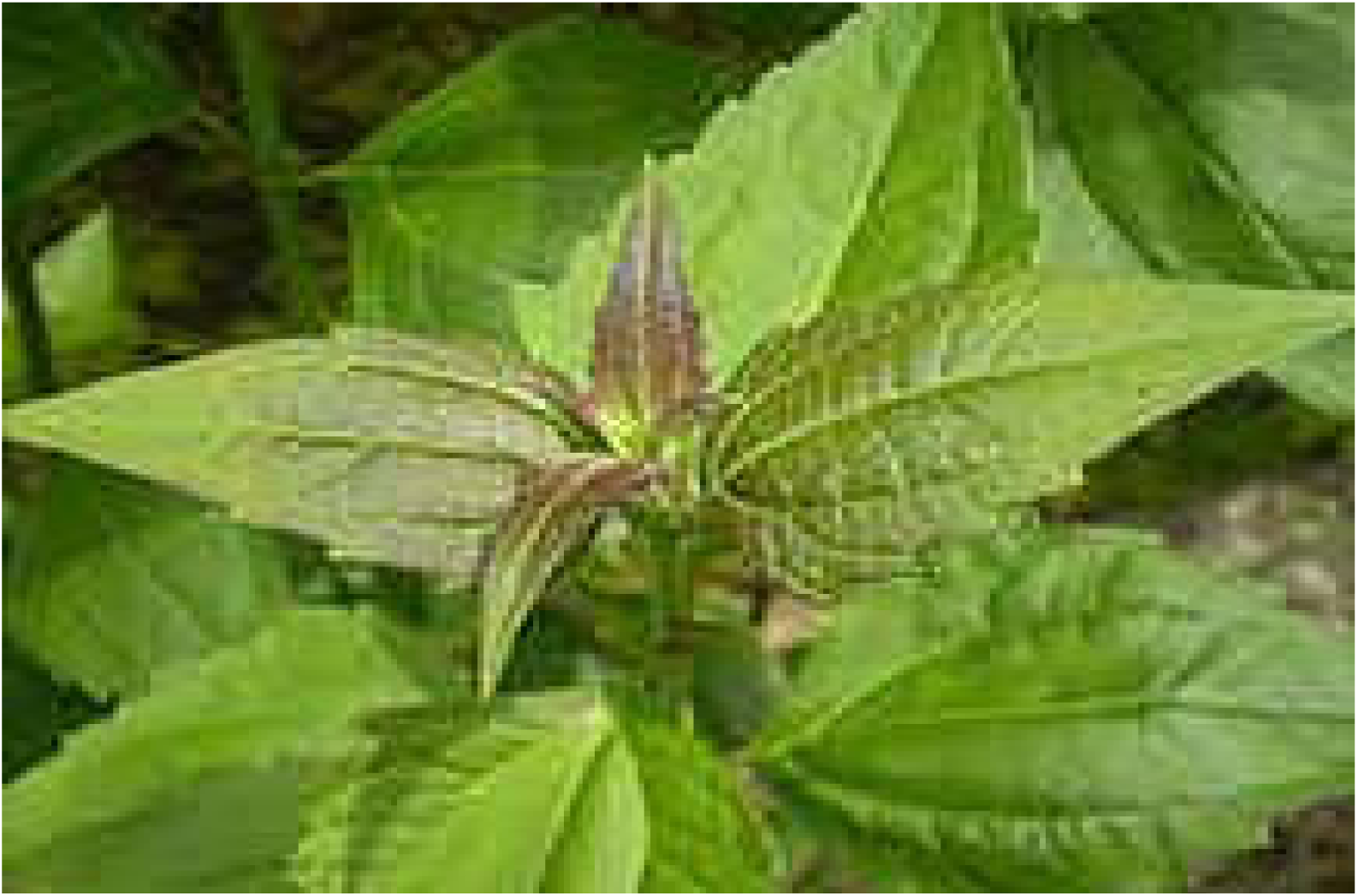
The leaves of Siam weed *Chloromolaena odorata*

